# A C++ library for protein sub-structure search

**DOI:** 10.1101/2020.04.26.062612

**Authors:** Jianfu Zhou, Gevorg Grigoryan

## Abstract

**Summary:** MASTER is a previously published algorithm for protein sub-structure search. Given a database of protein structures and a query structural motif, composed of multiple disjoint segments, it finds all sub-structures from the database that align onto the query to within a pre-specified backbone root-mean-square deviation. Here, we present an improved version of the algorithm, MASTER v.2, in the form of an open-source C++ Application Program Interface library, thereby providing programmatic access to structure search functionality. An entirely reorganized approach to database representation now enables large structural databases to be stored in memory, further simplifying development of automated search-based methods. Given the increasingly important role of structure-based data mining, our improved implementation should find ample uses in structural biology applications.

**Availability:** MASTER is available at https://grigoryanlab.org/master/master-v2.php.

## 1 Introduction

The apparent modularity of the protein structural universe (i.e., the frequent recurrence of local structural patterns) has been exploited in diverse structure modeling and design applications in a variety of ways (Grigoryan *et al*., 2011; Huang *et al*., 2014; Koga *et al*., 2012; Leaver-Fay *et al*., 2011; Szilagyi and Zhang, 2014). In our own work, we have found it particularly useful to identify precise atom-for-atom matches, from within the known structural database, to arbitrary constellations of disjoint backbone fragments (Frappier *et al*., 2019; Zheng and Grigoryan, 2017; Zhou *et al*., 2020). We previously developed the MASTER algorithm to address this computational task (Zhou and Grigoryan, 2015), and it has since been utilized by us and others (Kim *et al*., 2016; Lai *et al*., 2018; MacKenzie *et al*., 2016; Ojewole *et al*., 2018; Zhang *et al*., 2018; Zheng *et al*., 2015; Zheng and Grigoryan, 2017; Zhou *et al*., 2020). Here, we present the implementation of a MASTER Application Program Interface (API) library. It allows users to run MASTER search functions within the context of their own programs, making MASTER a more convenient tool for structural biology research.

## 2 MASTER API library

MASTER takes as query a structural fragment, composed of one or more disjoint segments, and provably finds all fragments from a database matching the query to within a given backbone root-mean-square deviation (RMSD) threshold (Zhou and Grigoryan, 2015). The method is fast, enabling searches over databases with tens of thousands of structures in a matter of seconds (for realistic thresholds), with the running time in practice most sensitive to the number of matches falling below the RMSD cutoff (Zhou and Grigoryan, 2015).

The previous implementation of MASTER comprised a stand-alone program that used an on-disk database. While it was possible to automate multiple search requests using system calls, the disk-based database meant considerable loss of efficiency due to I/O and inter-process communication. Here we present a new and improved implementation of MASTER in the form of an API library. It supplies MASTER search functions that mine a memory-accommodated database, abrogating the need for frequent disk I/O and allowing for the programmatic automation of search-based tasks without inter-process communication. To reduce the memory footprint of the database, we developed a proximity-search method and a data structure that eliminated the need for storing interatomic distances as in the previous implementation.

### 2.1 Room-saving proximity search algorithm

We refer as “proximity search” to the problem of identifying all points in a set that are within a given Euclidian distance of a query point. This is a common computational task of high utility in numerous areas. There exist several classes of approaches to this problem. An important class is based on space partitioning. For example, the traditional cell-list approach divides space into cells of identical size; points are mapped into these cells and proximity queries only happen within the same or neighboring cells (Frenkel and Smit, 2002). Another popular class of approaches is partitioning points themselves. For instance, in a bounding volume hierarchy, each point wrapped in a bounding volume forms the leaf node of a tree. Smaller bounding volumes are then clustered and wrapped within new larger ones in a recursive way, forming internal nodes of the tree, and the root encloses all the nodes using a single bounding volume in the end. Then, proximity queries performed between bounding volumes can efficiently prune sub-trees with no close points (Gottschalk, 2000). For a comprehensive review of representative methods adopted in applications related to molecular science, refer to (Artemova *et al*., 2011). In our new implementation of MASTER, we chose a simplified variant of the classical cell-list proximity search (Frenkel and Smit, 2002) described below.

As described previously (Zhou and Grigoryan, 2015), MASTER works by limiting the possible alignment locations of presently unaligned segments based on the deviations accumulated by the segments already aligned and the specified RMSD cutoff. To do so, MASTER relies on the ability to quickly perform residue proximity searches—i.e., to find the lists of residues located within specified ranges of distances from other residues. In our prior implementation of the algorithm, this functionality was supported via a database that explicitly stored, for every residue of every protein, a list of other residues from the same protein located within various distance bins of it. The relevant bins to check for a given proximity query could be identified from the desired inter-segment distance range (Zhou and Grigoryan, 2015). While allowing for fascicle residue proximity searches, this database design required *O*(*MN*^2^) of storage space (where *N* is the number of residues per protein and *M* is the number of proteins in a database), such that realistically-sized databases could not easily fit in memory on commodity machines.

Our new MASTER API implementation does not explicitly store interresidue distances and rather embeds every database structure into a 3D grid on the fly, upon reading. The minimal and maximal values of atomic coordinates along X-, Y-, and Z-axes define the vertices of the grid, and each residue (or rather its CA atom) is mapped into a grid box. Following this embedding, a proximity search simply involves identifying the box to which the source atom belongs followed by visiting only those boxes of the grid that could potentially contain atoms within the relevant distance range of the source atom.

A grid structure requires *O*(*n*^3^) space, where *n* is the linear dimensionality of the grid (i.e., number of boxes along each dimension). However, overly fine grids are not necessary in practice, since volume exclusion limits how closely protein atoms can be spaced, such that the physical grid box dimension should be roughly a constant. Suppose we choose grids with boxes of side length *l*. Then, to accommodate a protein with a radius of gyration *D*, we would need a grid with *n* = *O*(*D*/*l*) boxes along the side. For globular proteins, the radius of gyration should grow roughly as cubic root of length (i.e., number of residues), such that we would need 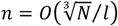 grid points along the side for a total memory cost of *O*(*N*/*l*^3^) = *O*(*N*). In the worst case of a fully extended protein, where the radius of gyration is proportional to length (i.e., *D* = *O*(*N*)), accommodating it in a grid would require *n* = *O*(*N*/*l*) grid points along the side or a memory cost of *O*(*N*^3^/*l*^3^) = *O*(*N*^3^). While the worst-case memory use is cubic in protein length, it does not represent a commonly encountered scenario. This is because in practice sub-structure queries are most meaningful when run with spatially local structural motifs—e.g., for the purpose of identifying dependences or preferences within them or to annotate a common functional unit. Because queries are expected to be well localized in space, it is not needed to store very long extended structures (even if these exist in the database) as single units. Rather, one would split such database entries into multiple overlapping portions, each more spatially localized (as a globular protein). Furthermore, radii of gyration for structures in the Protein Data Bank (PDB) do indeed appear to grow roughly as the cubic root of length (see Fig. 1), indicating that most entries are globular. Given all of this, *O*(*N*) is a better estimate of asymptotic memory usage in practice, which compares favorably to the *O*(*N*^2^) in our previous implementation (Zhou and Grigoryan, 2015).

**Fig. 1.**
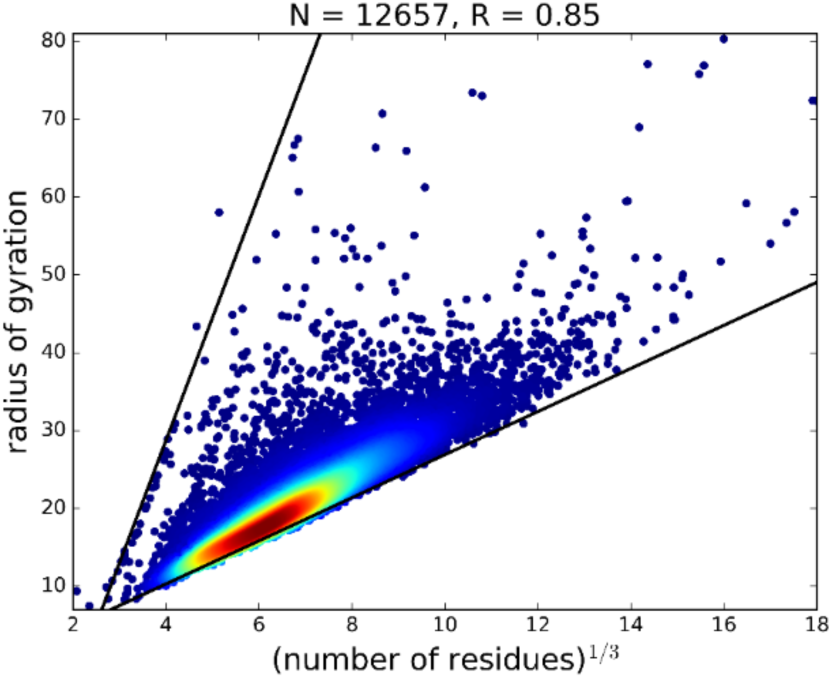
Cubic root of number of residues correlates with radius of gyration. Each point corresponds to a single protein structure in a nonredundant subset of the Protein Data Bank selected by using BLASTClust at 30% sequence identity (Altschul *et al*., 1990), with its radius of gyration plotted against the cubic root of its length. Point color indicates data density (decreasing order red-to-blue). Best-fit lines for the upper and lower boundaries, in solid black, are produced using least-squares fitting. The number of protein structures (**N**) and the Pearson correlation (**R**) are shown in the title.

The different data structure used in our new implementation impacts how a proximity search query is performed. Previously, MASTER explicitly stored the list of atoms within different distance bins (of width *r*) of each source atom. This took considerable memory but meant that all atoms within some distance *d* of a query atom could be found by visiting atom lists in *O*(*d*/*r*) = *O*(*d*) bins. In total, all atoms within distance of at most *d* + *r* were visited. If we assume some constant upper bound of atomic density *D*, this amounted to *O*(*D* · (*d* + *r*)^3^) = *O*(*d*^3^) atoms, such that the cost of a single proximity search was *O*(*d* + *d*^3^) = *O*(*d*^3^). In our new implementation, a proximity search is performed by visiting each grid box that is within the query distance *d* of the source atom. Assuming grid boxes with constant side length *l*, this means visiting *O*((2*d*)^3^/*l*^3^) = *O*(*d*^3^) boxes. The total number of atoms within these boxes is *O*((2*d*)^3^*D*) = *O*(*d*^3^) such that the overall cost of finding neighbors is *O*(*d*^3^ + *d*^3^) = *O*(*d*^3^). Thus, asymptotically, the running time of a proximity search remains the same between the two implementations. Further, in practice, once the disk I/O overhead associated with database reading is removed (substantially more costly in our old implementation), we found the running time of the two implementations to be comparable.

## 3 Example of using MASTER API library

The MASTER API library supplies the essential search function and several auxiliary functions for customizing a query and obtaining search results. Table 1 shows a simple excerpt from a hypothetical C++ program that uses these functions. The MASTER class serves as the main point of access to the search functionality, so the program first defines a MASTER object S. Through S, the program inputs the search query using `setQuery(queryPDB)`, where `queryPDB` is a path to a PDB file. Next, the code defines a target database by using the `addTargets(databaseList)` function, with `databaseList` being a path to a file with a list of PDB file paths denoting database entries. This function is responsible for uploading the entire database into memory, such that having issued this call once, the same MASTER object can be used for an arbitrary number of searches without needing disk access. The RMSD threshold is specified via `setRMSDCutoff(rmsdCut)` and finally the actual search is run by calling `search()` on the MASTER object. Results are returned by the function `solutions()` and stored in a `masterSolutions` object, which provides several functions for interrogating and manipulating resultant matches, including the ability to access structure and sequence information for each match. We also include a Python interface for the MASTER API library, which enables database reading, search, and match writing capabilities from Python.

**Table 1:**
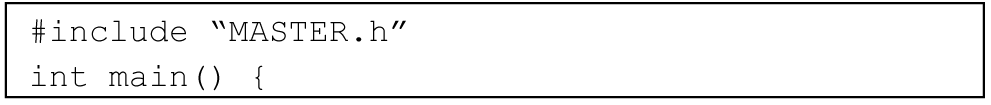
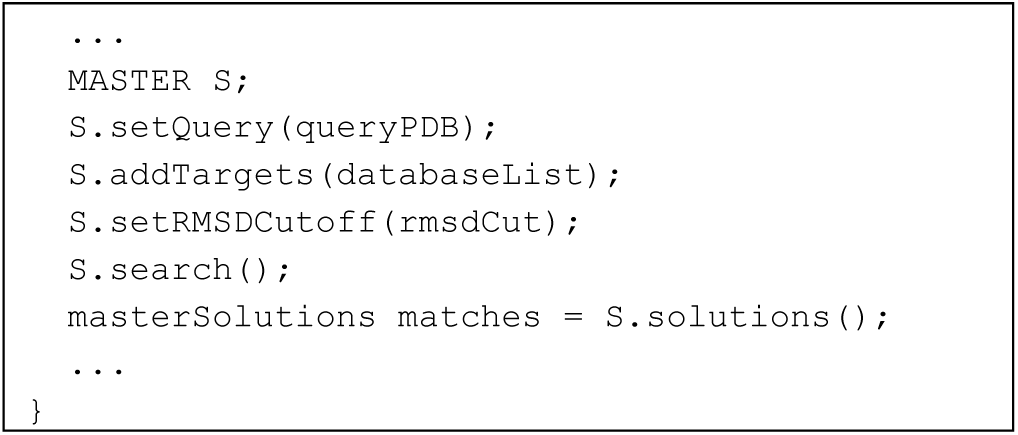
Example of using MASTER API library.

## 4 Conclusion

MASTER is a powerful tool for mining structural motifs in protein structural databases. Its API library provides researchers with a handy and flexible way to run MASTER searches in the context of their own applications.

## Funding

This work has been supported by NSF award DMR1534246 and NIH award P20-GM113132 (GG).

## Notes

### Competing Interest Statement

The authors have declared no competing interest.

